# The continuous differentiation of multiscale structural gradients from childhood to adolescence correlates with the maturation of cortical morphology and functional specialization

**DOI:** 10.1101/2024.06.14.598973

**Authors:** Yirong He, Debin Zeng, Qiongling Li, Lei Chu, Xiaoxi Dong, Xinyuan Liang, Lianglong Sun, Xuhong Liao, Tengda Zhao, Xiaodan Chen, Tianyuan Lei, Weiwei Men, Yanpei Wang, Daoyang Wang, Mingming Hu, Zhiying Pan, Haibo Zhang, Ningyu Liu, Shuping Tan, Jia-Hong Gao, Shaozheng Qin, Sha Tao, Qi Dong, Yong He, Shuyu Li

## Abstract

From childhood to adolescence, the structural organization of the human brain undergoes dynamic and regionally heterogeneous changes across multiple scales, from synaptic pruning to the reorganization of large-scale anatomical wiring. However, during this period, the developmental process of multiscale structural architecture, its association with cortical morphological changes, and its role in the maturation of functional organization remain largely unknown. Here, we utilized a longitudinal multimodal imaging dataset including 276 children aged 6 to 14 years to investigate the developmental process of multiscale cortical wiring. We used an in vivo model of cortical wiring that combines features of white matter tractography, cortico–cortical proximity, and microstructural similarity to construct a multiscale brain structural connectome. By employing the gradient mapping method, the gradient space derived from the multiscale structural connectome effectively recapitulated the sensory-association axis and anterior-posterior axis. Our findings revealed a continuous expansion of the multiscale structural gradient space during development, with the principal gradient increasingly distinguishing between primary and transmodal regions. This age-related differentiation coincided with regionally heterogeneous changes in cortical morphology. Furthermore, our study revealed that developmental changes in coupling between multiscale structural and functional connectivity were correlated with functional specialization refinement, as evidenced by changes in the participation coefficient. We also found that the differentiation of the principal multiscale structural gradient was associated with improved cognitive abilities, such as enhanced working memory and attention performance, and potentially supported by molecular processes related to synaptic functions. These findings advance our understanding of the intricate maturation process of brain structural organization and its implications for cognitive performance.

## Introduction

The human brain is a complex network that exhibits coordinated structural organizational principles at multiple spatial scales (1). From microscale neuron-to-neuron interactions to macroscale anatomical pathways connecting different brain regions, the anatomical connections encompass a range of scales (2, 3). Multiscale structural organization serves as a foundational framework to support various brain functions and is embedded within complex biological mechanisms (1, 2). Reconstructing the human brain structural connectome across multiple scales has implications for comprehending the principles of human brain organization and the foundation of cognitive function.

To comprehensively characterize neural organizations across multiple scales, an in vivo structural wiring model integrating complementary neuroimaging features based on multimodal magnetic resonance imaging (MRI) has recently been proposed (4). These features include macroscale structural characteristics, encompassing diffusion MRI tractography, cortical geodesic distance (GD), and microscale structural features called microstructural profile covariance (MPC) (4). By incorporating GD and microstructural similarity as additional structural connectivity features, this multiscale structural model compensates for the limitations of relying solely on white matter fibers as the predominant method for inferring structural connectivity (4). The GD quantifies the wiring cost and spatial proximity of the cortex (5). Moreover, the MPC evaluates the strength of structural connections by assessing microstructural similarity between cortical regions, based on the cortico-cortical “structural model”, which posits a close association between connectivity likelihood and similarity of cytoarchitecture across cortical regions (6, 7). Thus, these two features enhance the modeling of superficial and approximate distance connections within the gray matter (4). By employing the gradient mapping technique, previous studies revealed the existence of the principal organizational axis derived from the multiscale structural connectome in healthy adults (4) and individuals aged 14-25 years (8). Remarkably, this principal organizational axis spatially aligns with the principal axis of large-scale cortical organization known as the "sensorimotor-association (S-A) cortical axis" (9, 10). This axis signifies feature transition and functional processing across the cortical mantle from primary to association regions, capturing a hierarchical organization that manifests in anatomy (11), function(10), and evolution(12).

Childhood and adolescence (6-14 years of age) represent a critical period of rapid and continuous brain development marked by the restructuring of neural circuits influenced by puberty hormones. This restructuring leads to permanent brain structural reorganization and significant gains in cognitive and emotional functions, with a cognitive transition from concrete to abstract and logical thinking (13–15). Concurrently, the functional organization of the brain undergoes significant reconfigurations, with the principal axis shifting from a visual-sensorimotor gradient to a pattern gradient delineated by the S-A axis (16, 17). This period is also characterized by dynamic and regionally heterogeneous changes in brain structural features across multiple scales. For example, there are pronounced changes at the microscale level, including the growth of intracortical myelination and synaptic pruning (18, 19). Moreover, the maturation of white matter leads to a substantial reorganization of large-scale brain structural networks (20, 21). Consequently, delineating the development of multiscale structural organization during this period can yield structural insights into the significant functional reorganization and cognitive development.

From childhood to adolescence, cortical morphology undergoes remarkable refinements, including cortical surface area (SA) expansion and cortical thinning (22–24). Previous studies associated cortical morphology with multiscale structural connectivity, revealing that regions with similar morphological features were more likely to exhibit axonal connectivity and to share comparable cytoarchitecture (25, 26). In addition, biological processes potentially linked to the refinement of multiscale structural wiring architecture, such as microscale myelin proliferation into the periphery of the cortical neuropil, dynamic synapse reorganization, macroscale white matter fiber development, and axonal mechanical tension, are hypothesized to contribute to the maturation of cortical morphology (27–32). Thus, the potential association between the development of multiscale structural gradients and regionally heterogeneous maturation of cortical morphology warrants further exploration. Furthermore, although dynamic functional interactions between brain regions are constrained by invariant multiscale structural wiring, divergence between structural and functional networks may support flexible and diverse cognitive functions (1, 33). Corresponding to the development of structural brain networks, large-scale functional networks exhibit a shift toward a more segregated network topology, facilitating flexible and specialized brain functions (34–37). Therefore, it is worthwhile to investigate how structural constraints contribute to the maturation of functional organization and cognitive development. In addition, accumulating evidence indicates that genetic factors closely regulate the development of brain structure across regions (38). Axon guidance, which is closely linked to the formation of neural circuits during neural development, is associated with structural wiring (39–41). Therefore, investigating associated gene expression can reveal the underlying biological mechanisms driving multiscale structural development processes.

In this study, we utilized a longitudinal dataset of 437 scans, encompassing multimodal images from diffusion MRI (dMRI), T1-weighted (T1w) MRI, T2-weighted (T2w) MRI, and resting-state functional MRI (rs-fMRI), from 276 developing children (aged 6-14 years). Using the gradient mapping algorithm and linear mixed effect models, we first characterized the developmental patterns of multiscale structural gradients during childhood and adolescence. Furthermore, we explored the associations of these gradients with the refinement of cortical morphology. We also examined the associations between multiscale structure–function coupling and the maturation of cortical organization. Moreover, we investigated the underlying genetic basis and examined the relationships between multiscale structural gradients and individual cognition.

## Results

### Age-related changes in multiscale structural gradient during development revealed the gradual maturation of the S-A axis

We examined 437 scans, including structural MR, diffusion MR, T1w and T2w images, from 276 children aged 6-14 years (135 females) in a longitudinal dataset from the Children School Functions and Brain Development Project in China (Beijing Cohort) (CBD). To compute the multiscale structural gradients for each scan, we utilized a complementary model that integrated three cortical structural connectivity features (GD, MPC, and dMRI tractography) mapped onto a Schaefer 1000 parcellation (42). By implementing the diffusion map embedding algorithm, a set of components was derived and arranged in descending order based on the proportion of the variance accounted for by the component (Fig 1A, middle panel). We focused on the first two gradients, as they collectively accounted for a substantial proportion (approximately 45%) of the variance in cortical connectivity and represented principal axes of spatial variation in cortical wiring. Consistent with the two gradient patterns observed in previous studies of individuals aged 14-25 years and adults (4, 8), the principal gradient differed between the primary regions (somatomotor network [SN] and visual network [VN]) (positive values) and transmodal regions (default mode network [DMN]) (negative values), reflecting the hierarchical organization of the cortex. The second gradient demarcated the anterior and posterior cortex. To demonstrate the overall pattern of age-related changes in gradients, we computed group-averaged gradients for six age groups (6-7, 8, 9, 10, 11, and 12-13 years) and compared their global distributions. The group-averaged gradient maps for each group are shown in Supplementary S1 Fig. Our results demonstrated a consistent trend of the principal gradient becoming progressively distributed toward both ends during development (Fig 1B). Subsequently, we summarized the first two gradients at the network level according to intrinsic functional communities (43) and the atlas of laminar differentiation (44), as illustrated in Fig 1C. Our analysis demonstrated an increase of the principal gradient in the primary regions (SN, VN) and the dorsal attention network (DAN) and a decrease in higher-order networks, including the ventral attention network (VAN), limbic network (LN), frontoparietal network (FPN), and DMN. These findings also suggested an expansion pattern in the first gradient, which was further supported by results derived from the laminar differentiation atlas (Fig 1C, right panel). The second gradient showed an increase in the DAN and a decrease in the VN and LN throughout development (Supplementary S2A, B Fig). We next constructed a 2-dimensional gradient space to qualitatively assess global distribution patterns in the 6–7-year-old, 9-year-old, and 12–13-year-old groups, as depicted in Figure 1D. The gradient space demonstrated an expansion trend throughout development (the developmental process of the gradient space across different ages is depicted in Supplementary S1 Movie). Similar observations were also documented in the Schaefer 400 atlas (Supplementary S3 Fig and S2 Movie).

**Fig 1.**
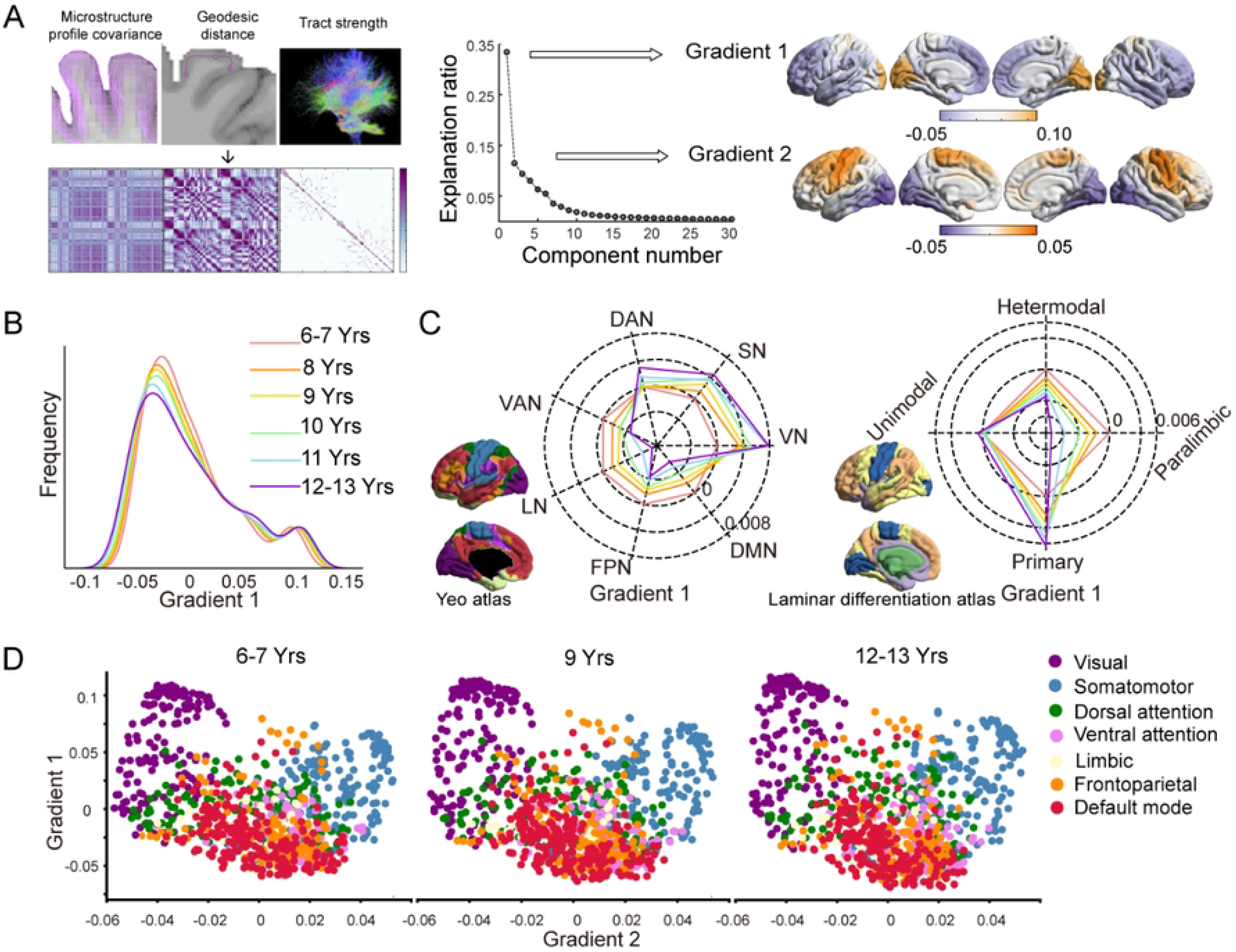
Multiscale structural gradients during childhood and adolescence. **(A)** The matrices containing the structural features of geodesic distance, microstructural profile covariance, and diffusion MRI tractography were concatenated and transformed into an affinity matrix, followed by the diffusion map embedding algorithm. The first two gradients capture the largest proportion of the variance. The group-averaged gradients were projected onto the cortical surface and visually represented (right). (**B)** The global density map of the principal gradient for six age-specific groups showed a gradual dispersal pattern with development. (**C)** Radar plot of the principal gradient for comparison between the 6–7-year-old group and other age-specific groups based on Yeo functional networks (left) (45) and laminar differentiation parcellation (right) (44). **(D)** The first two structural gradients mapped into a 2D gradient space for the 6–7-, 9-, and 12–13-year-old groups demonstrated an expansion pattern during development.

To quantify the effect of age on multiscale structural gradients during development, linear mixed-effect (LME) models (candidate models are described in Materials and Methods, Table 1) were constructed, and the optimal model was chosen based on the Akaike information criterion (AIC) (46). We first computed several global measures to describe the overall characteristics of the first two gradients, including the explanation ratio, range, and standard deviation. A higher explanation ratio signified a more prominent role in the organization of the structural connectome, while the range indicated differentiation between extremes, and the standard deviation measured inconsistency. We observed age-related increases in the principal gradient (age effect p < 0.001) and decreases in the second gradient (age effect p < 0.001) for all three global measures (Fig 2A). Additionally, dispersion was calculated by summing the Euclidean distances between each point and the centroid within the 2D space formed by the first two gradients for each individual, providing a quantification of the overall dissimilarity within the gradient space. The gradient dispersion exhibited an increasing pattern during development (Fig 2B). These findings indicated a shift toward a more distributed structural network topology during development, with the principal gradient increasingly differentiating between primary and transmodal regions. This finding is consistent with the increasing dominance of the principal gradient. In contrast, the second gradient suggested a progressive weakening of the anterior-posterior pattern.

**Fig 2.**
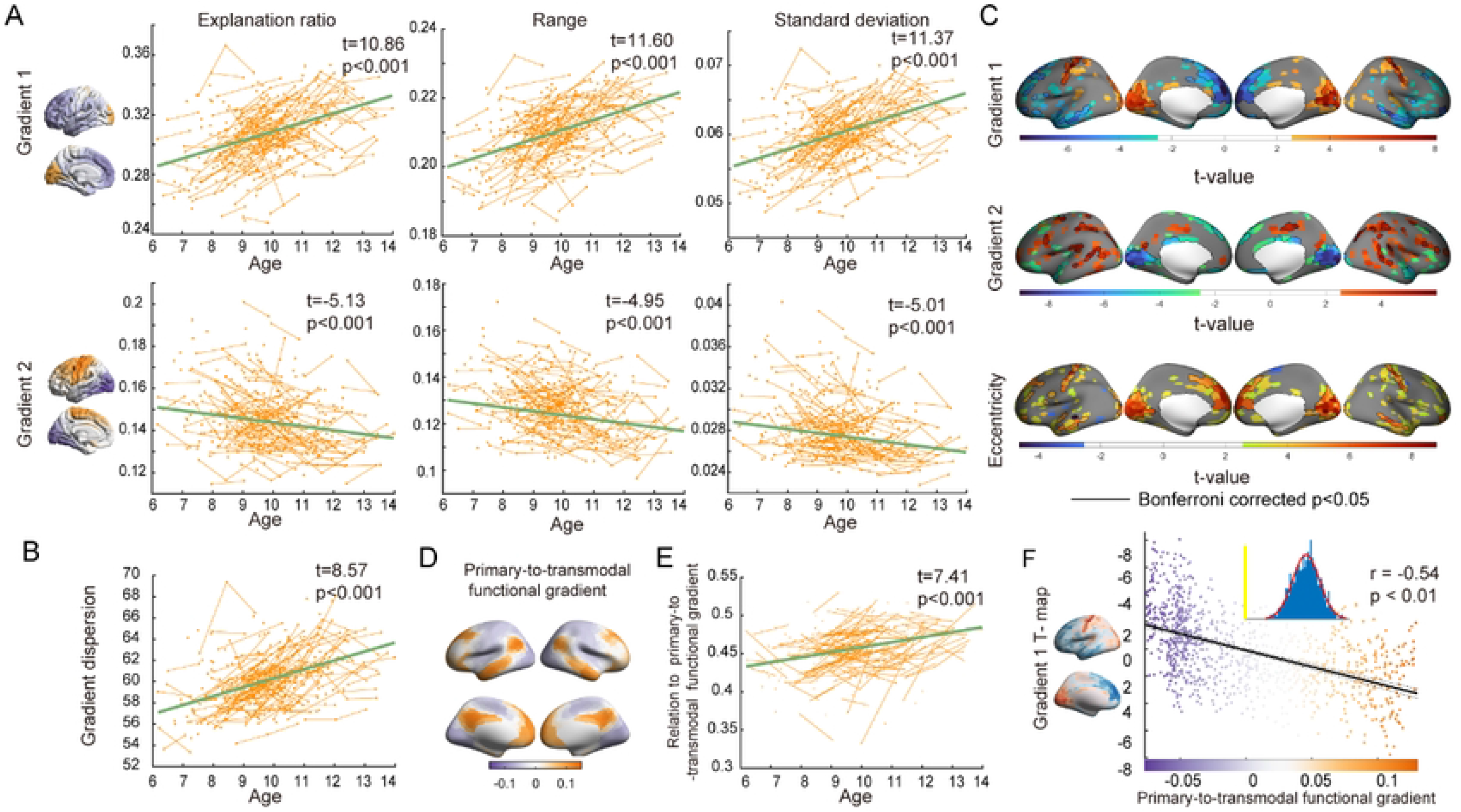
Age-related changes in gradients at both the global level and the node level. **(A)** Global measures of the first two gradients changed across age groups, including the explanation ratio (left), range (middle), and standard deviation (right) (age effect p < 0.01). **(B)** Age-related changes in gradient dispersion computed from the first two gradients (age effect p < 0.01). (**C)** T-statistic of age-related changes in nodewise gradients and the eccentricity map (p<0.05). The results that survived Bonferroni correction are circled by black lines (Bonferroni corrected p<0.05). **(D)** The primary-to-transmodal functional gradient derived from the group-averaged functional connectivity matrix. **(E)** Age-related changes in the correlation coefficient between the multiscale structural principal gradient and the primary-to-transmodal functional gradient (age effect p < 0.01). **(F)** Spatial correlation between the structural principal gradient age-related t-map and the primary-to-transmodal functional gradient. Each dot represents a brain node. The significance level was corrected for spatial autocorrelation (p _surrogate_<0.01).

**Table 1.**
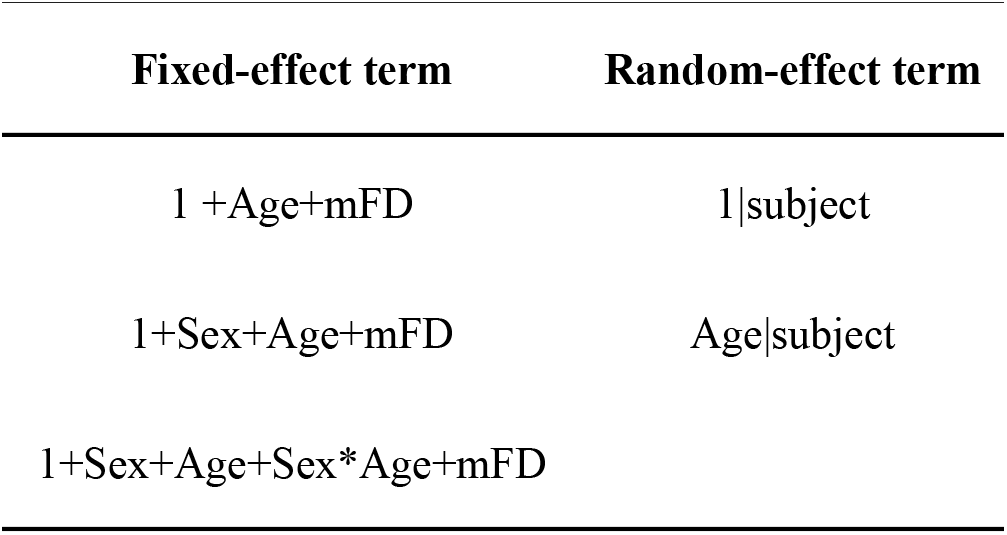
Candidate effects of mixed-effect models.

To examine the statistical age effect across the whole brain, we also leveraged the LME model at the node level. As depicted in Fig 2C, the principal gradient revealed age-related increases in the SN and VN corresponding to the positive extremum (t>1.976), while regions associated with the negative extremum (t<-1.967), such as the temporal, medial, and lateral prefrontal lobes, exhibited a pattern of decline (p < 0.05, Bonferroni corrected). For the second gradient, a significant decrease was observed in the VN (p < 0.05, Bonferroni corrected). Additionally, we calculated the eccentricity in each participant for each node by measuring the Euclidean distance between the given node and the centroid of the template gradient space derived from the averaged multiscale matrix. This metric quantified the deviation of each node from the central position. The eccentricity map demonstrated significant increases in the SN, VN, and medial lobe, corresponding to either end of the first gradient (p < 0.05, Bonferroni corrected). These results indicated an expansion of the gradient space during development, reflected in the strengthening differentiation of the principal gradient, which corresponded to the S-A axis. To further examine whether the S-A pattern of the principal gradient strengthened during development, we used the primary-to-transmodal functional gradient derived from the group-averaged functional connectivity (FC) matrix as the S-A axis (Fig 2D). We computed the correlation coefficient between the principal structural gradient and functional gradient for each scan. The LME model revealed a significant increase in the correlation coefficient during development, which indicated a strengthened S-A pattern in multiscale structural organization (t=7.41, p < 0.001) (Fig 2E). In contrast, the correlation coefficient between the second gradient and the functional gradient did not exhibit a significant effect of age (Supplementary S2C Fig). In addition, the age-related t-map of the multiscale structural principal gradient demonstrated a significant correlation with the primary-to-transmodal gradient, indicating temporal changes following the S-A organization pattern (r= −0.54, p_surrogate_<0.01) (Fig 2F). Therefore, this analysis demonstrated that multiscale structural wiring architecture shifted toward a more distributed hierarchical organization during childhood and adolescence.

### The multiscale structural principal gradient and its maturation are associated with the development of cortical morphology

Considering that cortical regions with similar morphological features are more likely to have structural connections and that structural connectivity features such as myelin and white matter tracts are potentially interrelated with the maturation processes of cortical morphology, we hypothesized that the refinement of the multiscale structural principal gradient may coincide with the heterogeneous maturation of cortical morphology. Subsequently, we employed five cortical morphometric measures that are relevant to the aforementioned biological processes and delineated a comprehensive cortical morphological profile. These measures included cortical thickness (CT), gray matter volume (GMV), SA, mean curvature (MC), and Gaussian curvature (GC). We investigated the associations between the multiscale structural principal gradient and morphometric features (Fig 3A). Given the similarities in the spatial patterns of these metrics, we performed principal component analysis (PCA) to project the five features onto a set of principal axes that effectively captured the spatial variation in the cortical morphological profile. The first component (PC1) explained nearly 85% of the variance, and we incorporated PC1 into subsequent analyses. As shown in Fig 3B, the group-averaged PC1 exhibited differentiation between primary regions (i.e., the SN and VN) and transmodal regions (i.e., the FPN and DMN), indicating that distinct morphometric attributes distinguish these two types of brain regions. Then, as depicted in Fig 3C, we explored the relationship between PC1 and multiscale structural gradient 1 and identified a strong correlation (r=0.69, p_surrogate_ <0.01). These findings suggested a potential association between cortical morphology and cortical wiring architecture across the cortical mantle, as regions exhibiting similar morphological features also display comparable multiscale structural connectivity profiles.

**Fig 3.**
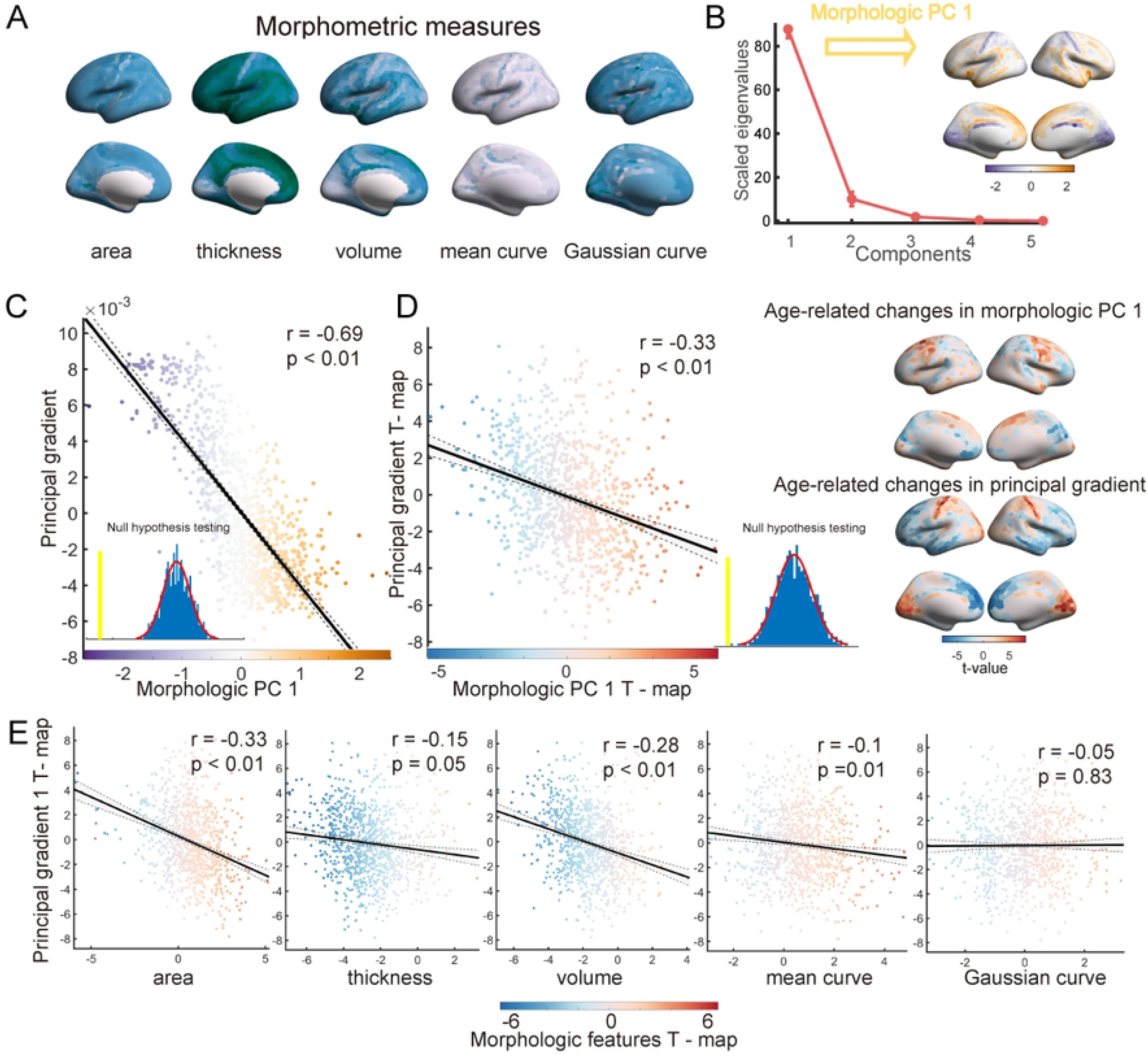
Association between the multiscale structural principal gradient and morphometric features. **(A)** Group-averaged morphometric features, including cortical thickness, gray matter volume, surface area, mean curvature, and Gaussian curvature. (**B)** The five morphometric features were input into the PCA algorithm, and components were ordered according to the proportion of variance they accounted for. The principal component (PC1) was mapped on the surface (right). **(C)** Spatial correlation between the multiscale structural principal gradient and morphometric PC1. Each dot represents a brain node. The significance level was corrected for spatial autocorrelation (p _surrogate_<0.01). **(D)** Spatial correlation between age-related t-maps of the multiscale structural principal gradient and morphometric PC1 (p _surrogate_<0.01). **(E)** Spatial correlation between age-related t-maps of the multiscale structural principal gradient and morphometric features, including surface area, cortical thickness, gray matter volume, mean curvature, and Gaussian curvature.

To validate the presence of a developmental association between cortical wiring and cortical morphology, we investigated the spatial correlation of mature patterns between them. Specifically, we employed the previously mentioned LME model on PC1 to characterize the effect of age on cortical morphology. As illustrated in the right panel of Fig 3D, we observed an increase in the prefrontal lobe, which occupies the positive end of PC1. This observation suggested distinct maturation processes between the prefrontal lobe and other brain regions. Moreover, as shown in the left panel of Fig 3D, the correlation analysis between the t-maps of multiscale structural gradient 1 and morphometric PC1 revealed a congruent developmental pattern with a correlation coefficient of r=-0.33 (p_surrogate_ <0.01). The increase in the multiscale structural principal gradient in the SN was accompanied by a decrease in PC1, while the decrease in the principal gradient in the prefrontal and temporal lobes was accompanied by an increase in PC1. The obtained results validated our hypothesis that there are synchronized maturation patterns between cortical wiring and cortical morphology. As shown in Fig 3E, to investigate the extent to which individual morphological features co-evolve with the multiscale structural gradient, we also conducted a correlation analysis between the t-map of multiscale structural gradient 1 and the t-map of each morphometric feature. Notably, a significant association was observed between t-maps of the principal gradient and SA (p_surrogate_ <0.01), GMV (p_surrogate_ <0.01), and MC (p_surrogate_ = 0.01). These findings provide evidence of interconnected spatial patterns and developmental influences between the multiscale structural connectome and cortical morphology.

### Development of multiscale structure–function coupling associated with the refinement of cortical functional specialization

The coupling between structure and function indicates that structure is the fundamental framework that facilitates synchronized fluctuations in functional activities underlying cognition (47). To further investigate the role of the multiscale structural connectome in shaping the development of functional architecture, we analyzed the coupling between structure and function for each region. Coupling was assessed through Spearman rank correlation between the connectivity profiles of structure and function (Fig 4A). As shown in Fig 4B, the group-averaged coupling map revealed distinct patterns across the cortex, ranging from −0.01 to 0.34, reflecting the alignment of functional and multiscale structural connectivity profiles of the given region. The network-level analysis, based on intrinsic functional communities (43), further revealed a hierarchical pattern across the cortical mantle characterized by greater levels of coupling in primary regions and lower levels in transmodal regions (Fig 4C). A previous study revealed that variability in structure‒function coupling is related to functional specialization (47). To investigate whether multiscale structure‒function coupling is associated with functional specialization, we calculated the participation coefficient (PaC) for each node based on both multiscale structural and functional networks. The PaC was employed to assess intermodule connectivity and quantify the degree of each node’s involvement in other functionally specialized modules. Nodes with lower values indicated a greater degree of functional specialization. The correlation between multiscale structure‒function coupling and group-averaged PaC maps is illustrated in Fig 4D, revealing a significant relationship (correlation with structural PaC: r = −0.61, p_surrogate_ <0.01; functional PaC: r = −0.51, p_surrogate_ <0.01). These findings indicated that greater structure‒function coupling was associated with greater functional specialization, while lower coupling corresponded to greater functional integration. Furthermore, we demonstrated that structure–function coupling aligned with both structural and functional hierarchies (correlations with the multiscale structural gradient: r = 0.39, p_surrogate_ <0.01; functional gradient: r = −0.55, p_surrogate_ <0.01) (Supplementary S4A, B Fig). These findings demonstrated that the coupling of multiscale structure and function reflected functional specialization and hierarchy.

**Fig 4.**
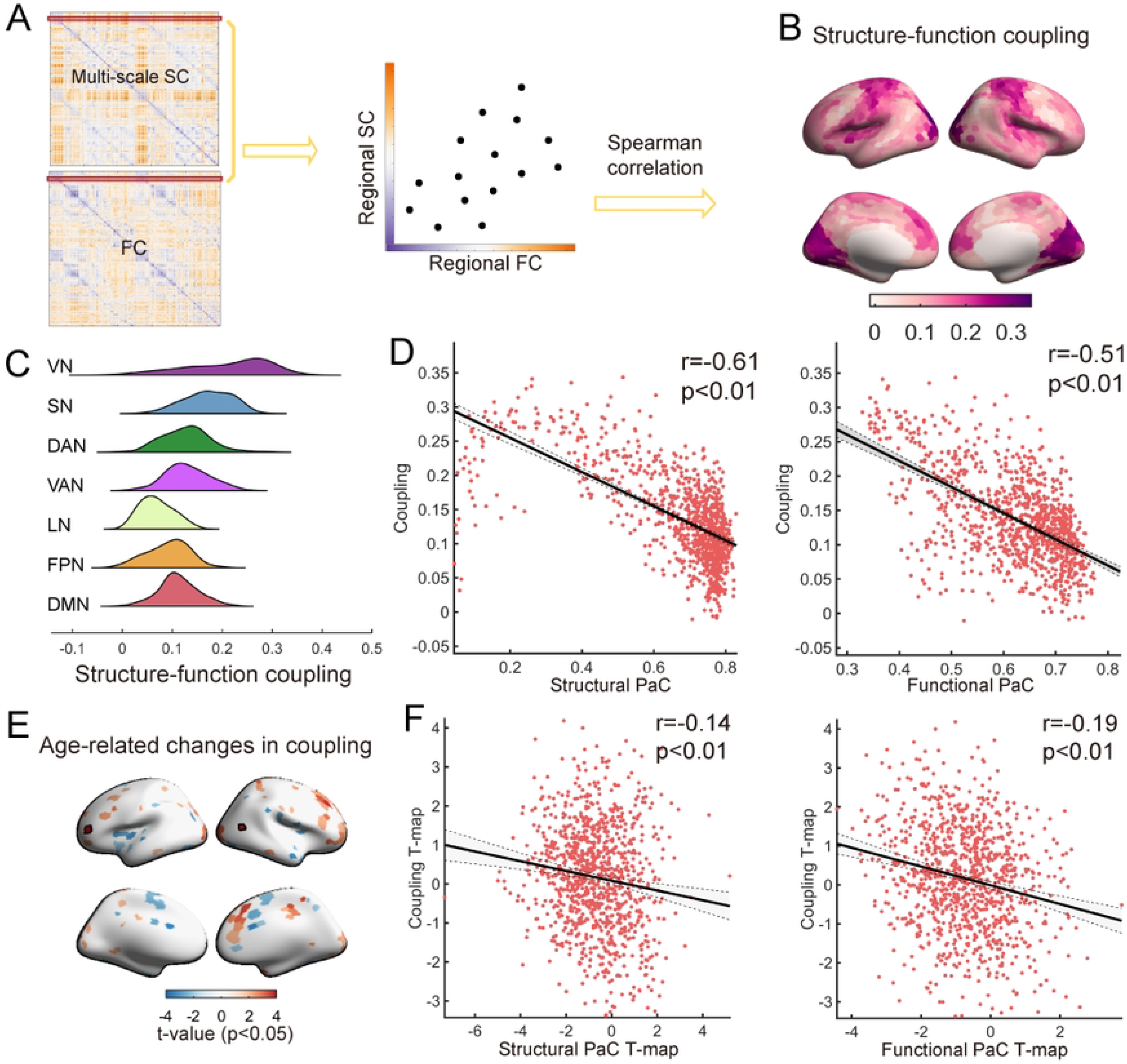
Multiscale structure–function coupling during development. **(A)** For each region, multiscale structure‒function coupling was calculated as the Spearman correlation coefficient between the multiscale SC and FC profiles of that region. (**B)** A group-averaged multiscale structure–function coupling map of the cortical surface is depicted. **(C)** The distributions of the coupling map in Yeo functional networks (45). **(D)** Spatial correlation between the multiscale structure‒function coupling map and the structural/functional participation coefficient map (p _surrogate_<0.01). **(E)** Age-related changes in multiscale structure–function coupling. Age-related increases/decreases are shown in red/blue, and the results surviving false discovery rate (FDR) correction are circled by black lines. **(F)** Spatial correlation between t-maps of multiscale structure–function coupling and the structural/functional participation coefficients (p _surrogate_<0.01).

To characterize age-related changes in regional multiscale structure‒function coupling, we used the LME model. As depicted in Figure 4E, the prefrontal cortex exhibited enhanced coupling during development, whereas the insula demonstrated reduced coupling. The Yeo atlas was subsequently employed to provide a network-level summary of these findings; however, no statistically significant results were observed in the network-level analysis (Supplementary S5 Fig). Considering the close interplay between structure–function coupling and segregation, we further hypothesized that age-related changes in coupling are accompanied by alterations in the PaC. As depicted in Figure 4F, the correlation analysis between t-maps of coupling and structural as well as functional PaCs revealed a congruent developmental pattern (correlation with structural PaC: r = −0.14, p_surrogate_ <0.01; functional PaC: r = −0.19, p_surrogate_ <0.01). This finding suggested that brain regions exhibiting increases in structure‒function coupling were more likely to be accompanied by an increased degree of functional specialization. Taken together, these findings demonstrated that the maturation of multiscale structure‒function coupling was related to the refinement of functional specialization from childhood to adolescence.

### The differentiation of the principal multiscale structural gradient was related to better cognitive performance

Structural connectivity serves as the fundamental basis for neuronal interactions that underlie the emergence of cognition and behavior (33). Throughout childhood and adolescence, attention and executive function undergo continuous enhancement (48). Subsequently, we sought to explore the implications of cortical wiring for individual cognition by investigating two cognitive dimensions: working memory (WM) and attentional ability. WM is associated with complex tasks such as temporary storage and manipulation of information (49). Attention involves prioritizing task-relevant information processing while disregarding irrelevant information (49). Here, WM was measured by a typical numerical n-back task, while attention performance was measured by response time for alerting, orienting and executive control (EC) tasks (see Methods for further details). We next assessed the associations between the gradient data and cognition data across individuals via partial least square correlation (PLSC) analysis. PLSC offers a multivariate perspective that can capture complex relationships within multidimensional data. Considering the distinct cognitive aspects assessed by the two tests, separate PLSC analyses were performed for each cognitive domain. Through PLSC, we generated latent components (LCs) that captured the optimal associations between the principal gradient and cognitive scores.

For WM, the first LC (LC1) exhibited significance in the permutation test (p<0.01). For LC1, the composite scores were computed by projecting the original data onto their corresponding weights. The correlation between the WM composite score and the gradient 1 composite score was significant, indicating a strong positive relationship between the cognitive and gradient data (r=0.48, p<0.01) (Fig 5A). Additionally, we calculated the loadings of gradient 1 and WM by computing the Pearson correlation between the original data and the composite scores, thereby quantifying the contribution of the given brain (cognitive) measure for the LC. As shown in Fig 5B and Fig 5C, higher WM composite scores were associated with worse WM performance, while greater gradient composite scores were linked to higher values of gradient 1 in transmodal regions and lower values in primary regions. These significant loadings, tested by bootstrap resampling (n=1000), are depicted with shadows in WM and black lines in gradient 1. Better WM performance was associated with higher gradient 1 values in primary regions and lower values in transmodal regions.

**Fig 5.**
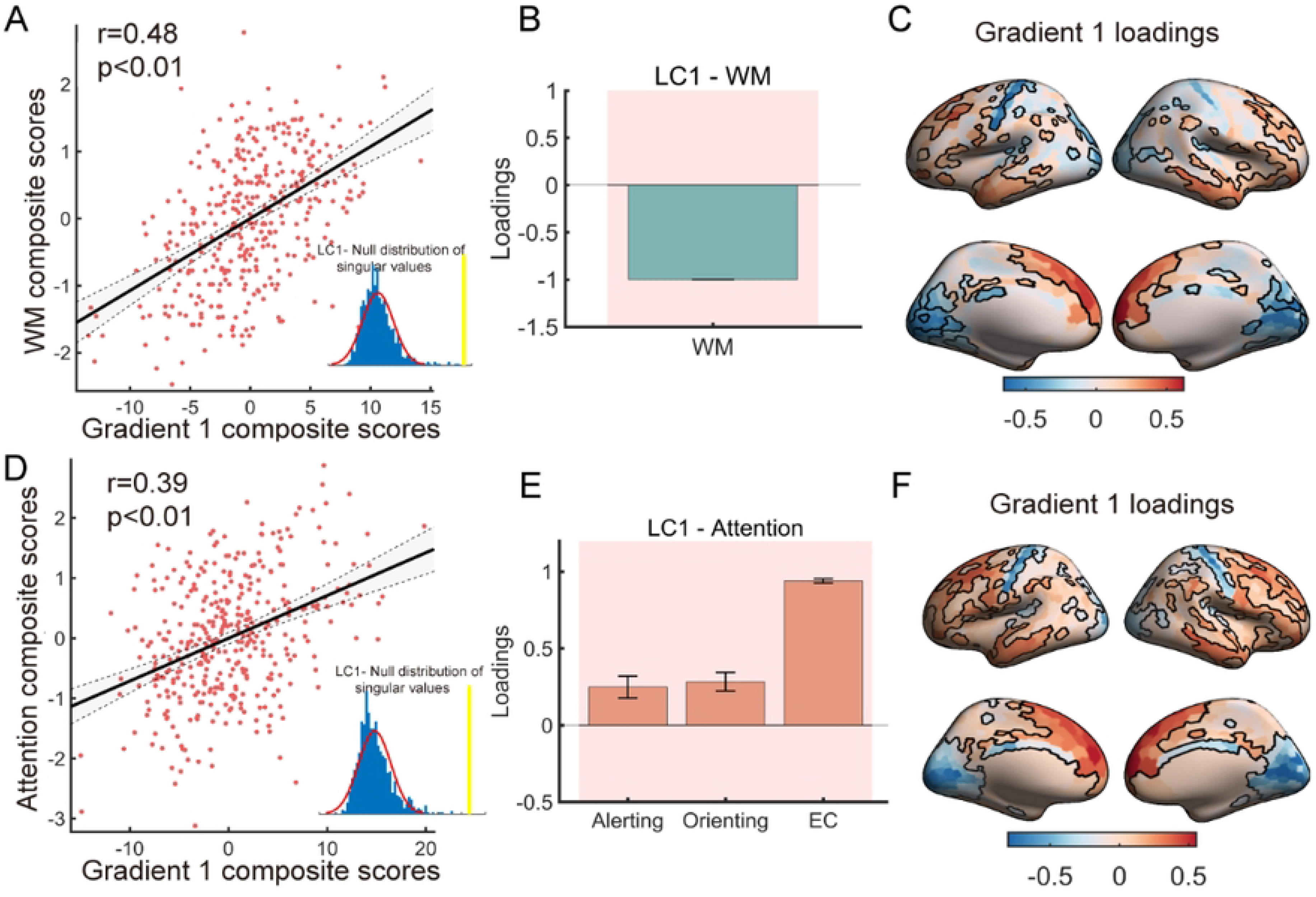
Partial least square correlation (PLSC) analysis revealed an association between the principal gradient and cognitive scores. **(A, D)** Pearson correlations between the principal gradient and composite scores of working memory/attention. The inset figure shows the null distribution of singular values estimated by the permutation test (n = 1000). (**B, E)** Loadings of WM/attention were calculated by Pearson correlation between the cognitive measurements and their composite scores. The shadows represent significant loadings tested by bootstrap resampling (n=1000). **(C, F)** Gradient loadings were calculated by Pearson correlation between gradient 1 and their composite scores. The loadings of regions with black lines were subjected to a significance test by bootstrap resampling (n=1000).

Similar to the WM results, LC1 derived from the attention-related PLSC analysis accounted for 46.57% of the covariance (p=0.001), showing a significant association between attention and gradient 1 composite scores (r=0.39, p<0.01) (Fig 5D). As shown in Fig 5E and Fig 5F, better attention scores were associated with higher gradient 1 values in transmodal regions and lower values in primary regions. Given that attention performance was measured through response time, larger attention scores indicated poorer attention performance. Therefore, these findings were consistent with the results obtained from the WM analysis, suggesting a significant association between improved cognitive performance and decreased negative value as well as increased positive value of the principal gradient (strengthened S-A pattern in multiscale structural organization). Consequently, these collective outcomes provide evidence that the enhancement of the S-A axis pattern along multiscale structural gradient 1 was associated with better cognitive performance.

### The maturation of the principal multiscale structural gradient was associated with gene expression profiles

To explore the underlying biological mechanisms of the maturation of multiscale structural gradients, we applied genome expression data from the Allen Human Brain Atlas (AHBA) (https://human.brain-map.org (50)). The microarray data were preprocessed using the abagen toolbox (version 0.1.3; https://github.com/rmarkello/abagen). Given that data from the right hemisphere were incomplete, we only used the data from the left hemisphere. By mapping the microarray data to the Schaefer 1000 atlas, we obtained a 416 ×15631 (region × gene) matrix (Fig 6A).

**Fig 6.**
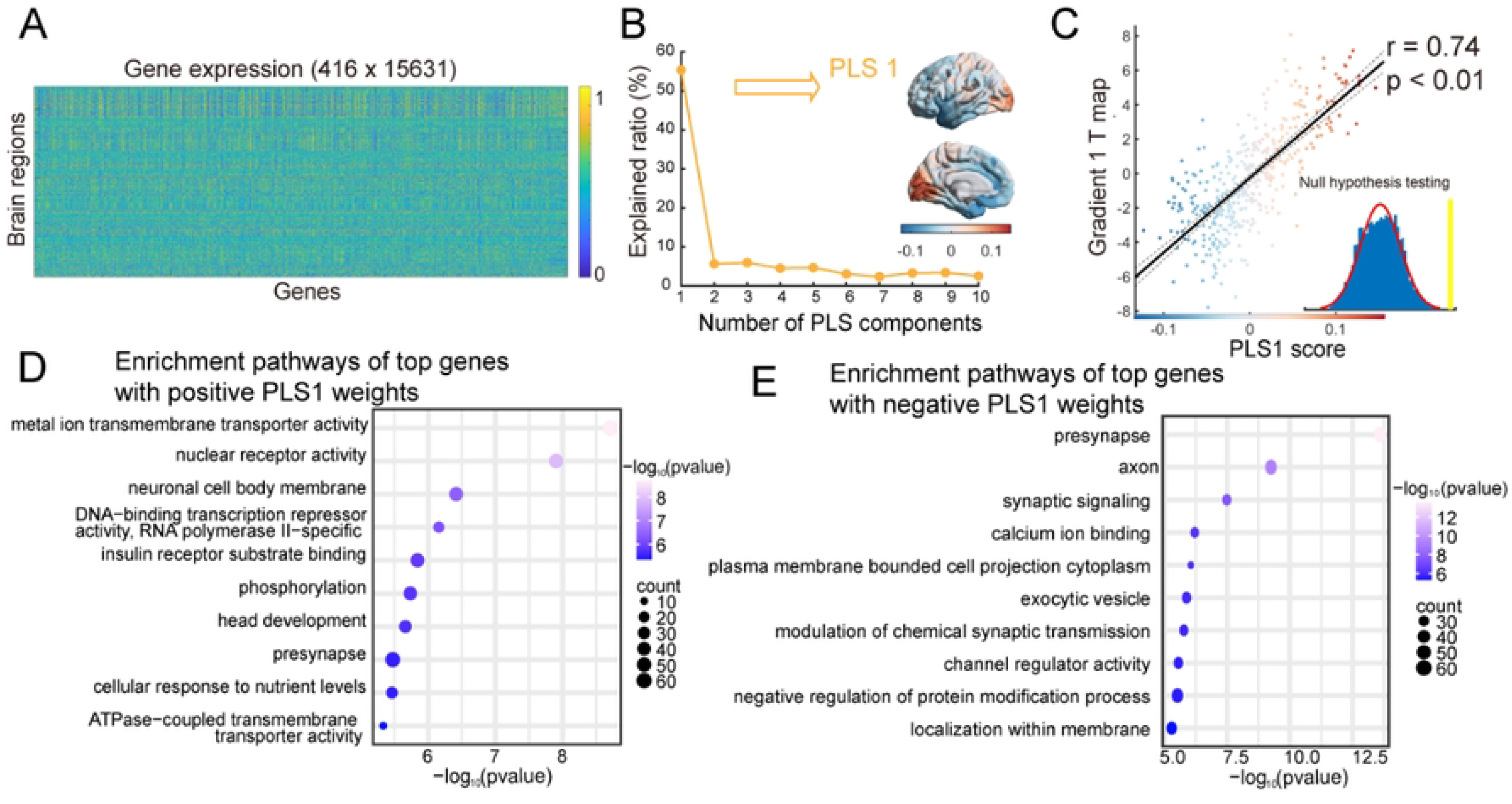
Association between age-related changes in the principal gradient and gene expression profiles. **(A)** Gene expression profiles across 416 brain regions. (**B)** The explained ratios for the first 10 components derived from the partial least squares regression algorithm. The first component (PLS1) accounted for the largest proportion of the variance and is depicted in the right panel. **(C)** Spatial correlation between age-related changes in the multiscale structural principal gradient and PLS1 scores. Each dot represents a brain node. The significance level was corrected for spatial autocorrelation (p _surrogate_<0.01). **(D, E)** Gene Ontology (GO) enrichment pathways of the top 10% of genes with positive/negative PLS1 weights. The 10 most significant GO terms are displayed (false discovery ratio-corrected).

Subsequently, we employed a partial least squares (PLS) regression algorithm to investigate the relationships between the age-related gradient 1 t-map and the gene expression matrix. The first component (PLS1) accounted for the largest proportion of the variance (55.35%) and represented the optimally weighted linear combinations of gene expression patterns (Fig 6B). The spatial pattern of PLS1 was spatially correlated with the multiscale structural gradient 1 t-map (r=0.74, p_surrogate_<0.01, corrected for spatial autocorrelation) (Fig 6C).

To further investigate the biological implications, the genes were ranked based on the weights from PLS1, and the top 10% of genes from both the positive (PLS1 +) and negative (PLS1 -) weights were input into the Metascape web tool for gene enrichment analysis and visualization (all p_FDR_<0.05) (51). Notably, the expression of positively weighted genes was positively correlated with the gradient 1 t-map.

Gene Ontology (GO) analysis was performed to identify related molecular functions, biological processes, and cellular components. As shown in Fig 6D, several meaningful brain development-related terms emerged for the PLS1+ genes, such as “head development”, “metal ion transmembrane transporter activity”, “neuronal cell body membrane” and “presynapse” (Fig 6D). On the other hand, the PLS1-genes were enriched in several synapse-related terms, such as “presynapse”, “axon”, “synaptic signaling”, “exocytic vesicle”, “modulation of chemical synaptic transmission”, and “calcium ion binding” (Fig 6E). The 20 most significant GO terms are depicted in Supplementary S6 Fig.

## Discussion

In this study, we documented the typical development process of multiscale structural gradients from childhood to adolescence based on an advanced structural connectome model. The results demonstrated that the maturation of a multiscale structural gradient was differentiated along the S-A cortical axis during the developmental period of 6-14 years of age. The shared developmental consequences of the multiscale structural gradient and cortical macrostructure indicated a potential interconnected maturation mechanism between the structural connectome and cortical morphology. The developmental changes in multiscale structure–function coupling reflected the refinement of functional specialization. In addition, the enhancement of the S-A axis pattern along the principal gradient demonstrated associations with enhanced cognitive performance and synapse-related gene expression. These findings provide a comprehensive understanding of the maturation principles of multiscale structural organization in the human brain during childhood and adolescence, as well as the underlying biological mechanisms involved.

### Differentiation of the multiscale structural principal gradient with development

The multiscale structural connectome model in this study integrated three complementary neuroimaging features, diffusion MRI tractography, MPC, and cortical GD (4). Tract strength is the dominant measure for assessing white matter connectivity, while GD can infer short adjacent cortico-cortical connections (4, 5). MPC measures similarities between cortical regions, as connectivity is more likely to exist between regions with similar cytoarchitectures (6, 7). Consistent with findings in healthy adults and adolescents aged 14-25 years (4, 8), our study identified two principal axes of multiscale structural connectome organization, the primary-transmodal axis and anterior-posterior axis, in an accelerated longitudinal cohort aged 6-14 years. In this population, both qualitative (Fig 1D) and quantitative (Fig 2B, C) analyses indicated an expanding gradient space during development that was mainly driven by the continuous differentiation of the principal gradient. Furthermore, given the more pronounced differentiation of the S-A axis, a primary-to-transmodal functional gradient was utilized as a proxy for this axis, and a tendency for the principal multiscale structural gradient to align with the S-A axis during development was revealed (Fig 2E). The continuous differentiation between the primary and transmodal cortex along the principal gradient aligned with the neurodevelopmental hierarchy from multiple findings, which suggested a varied developmental pattern between the primary and transmodal cortex (9). First, this differentiation pattern along the principal structural gradient mirrored the increasing differentiation across the functional hierarchy during this period, as indicated by the shift in the principal functional gradient from the visual-sensorimotor gradient toward a pattern gradient characterized by the S-A axis (16, 17). Second, this differentiation pattern was also consistent with evidence from white matter connectivity and myeloarchitecture, which demonstrated augmented differentiation of this axis during development (52, 53). In addition, the differentiation of cortical features along the S-A axis may delineate distinct cognitive functions and facilitate executive, socioemotional, and mentalizing functions within the transmodal region (9). Recent studies have indicated that differentiation along the S-A axis is related to flexible cognitive processing and better cognitive function (54, 55). Our results corroborated this finding that a more differentiated gradient along the S-A axis was related to better WM and attention performance.

### Interactions between the development of the multiscale structural gradient and cortical morphometric features

Our findings revealed coordinated spatiotemporal developmental patterns of cortical morphometric profiles that encompass multiple morphometric features and the principal multiscale structural gradient incorporating white matter and cortical microstructure. Some empirical evidence and theoretical hypotheses have established associations between changes in cortical morphology and structural wiring; one hypothesis is Seldon’s "balloon model" (56), which states that akin to an expanding balloon, the growth of white matter induces tangential stretching and thinning of its connected cortex. This hypothesis was supported by correlations found between cortical surface expansion and increased subcortical white matter fibers during development (30). The theory proposed by Essen (31) links the cortical folding pattern to axonal mechanical tension, with gyri potentially formed through mechanical tension pulling closely interconnected regions together. Gray matter thinning during childhood and adolescence is attributed to biological processes such as synaptic pruning, apoptosis (28, 57), and proliferation of myelin at the interface between gray matter and white matter (27–29). Previous studies also revealed associations between cortical thinning and increased white matter fibers during development (58, 59). Furthermore, considering the brain’s organization as a network of interconnected regions, a recent study adopting a network perspective demonstrated the constraints of the WM network on the maturation of CT from childhood to adolescence (60). Our study also revealed that regions exhibiting analogous structural connection profiles demonstrated congruent cortical morphology in spatial and maturation patterns, which can be elucidated through various mechanisms. First, structurally interconnected regions tend to possess similar cytoarchitecture and may develop during comparable time windows (61–63). Regions with similar cytoarchitectonic patterns tend to exhibit similar morphological characteristics (25). Second, the regionally heterogeneous developmental patterns of cortical morphology may be attributed to mutual trophic influences supported by structural wiring (64). Third, a recent study demonstrated that regions with similar cytoarchitectonic features and white matter interconnections are more likely to exhibit similar neurotransmitter receptor profiles (65). Consequently, these regions may be subject to coregulation through similar physiological mechanisms (60, 66). The findings of this study offer novel insights into the interconnected maturation mechanisms between cortical wiring and macrostructure, suggesting a potential role for structural connectivity in shaping cortical morphology.

### Relationships between changes in multiscale structural organization and functional organization during development

Our study revealed a continuous differentiation pattern along the principal multiscale structural gradient during development, paralleling the primary-to-transmodal functional gradient results reported by (17) in the same population as ours. This finding indicated a harmonized process of structural and functional maturation in human brain development, characterized by increasingly enhanced hierarchical organization and segregated topology. Previous studies also highlighted the synchronized maturation of structural and functional organization. A study based on functional intrinsic cortical activity revealed a hierarchical neurodevelopmental axis, which was linked to a progressive increase in intracortical myelination (67). Moreover, throughout the developmental process, both the structural and functional topology displayed a more distributed and segregated pattern (68, 69). These results suggested a mature process of enhanced segregation, manifested in structural and functional synchronization.

In addition, numerous studies have consistently demonstrated that structure‒function coupling exhibits regional heterogeneity, with the degree of coupling aligning along the S-A axis (47, 70, 71). Our findings supported the prevailing trend, with a greater degree of coupling in the primary cortex than in the transmodal cortex. The primary regions exhibit more rapid and accurate responses to external stimuli, necessitating stronger structural constraints. In contrast, the transmodal regions are untethered from structural constraints, consistent with their more flexible and diverse functional roles (6, 72). Low coupling in the transmodal cortex may be related to functional flexibility and diverse task demands (73). A previous study utilizing the white matter connectivity network and functional network demonstrated that coupling reflects functional segregation (47). Consistent with this study, our study also revealed a significant spatial correlation between multiscale structure–function coupling and the PaC, as well as their interrelated developmental patterns. Our findings revealed that during development, regions exhibiting stronger coupling between structure and function demonstrated stronger functional specialization, characterized by a greater degree of segregation. Conversely, regions with weaker coupling showed a greater degree of integration. Notably, stronger coupling between structure and function supports faster and more accurate specialized functions, while regions with fewer structural constraints are associated with greater flexibility and integrative roles (6, 72). These results established a compelling connection between structural-functional coupling and the underlying mechanisms of cortical organization.

### Transcriptional profiling of the developmental multiscale structural gradient

Using gene expression data from the AHBA dataset, our transcriptome analysis revealed that developmental changes in multiscale structural gradient 1 were associated with the transcriptional profiles of genes involved in development- and synapse-related terms, such as “presynapse”, “axon”, “synaptic signaling”, and “calcium ion binding”. Synapses serve as the foundation for communication between neurons in the nervous system. The elimination of synapses persists throughout development, with the pruning process exhibiting heterogeneity across brain regions and refining functional circuits (19, 74). Sensory regions complete this process during late childhood, while higher-order regions continue to experience synaptic pruning into adolescence (75). Calcium ions trigger the release of neurotransmitters and initiate synaptic transmission (76). Myelinated axons serve as the primary conduits for transmitting information within the central nervous system, constituting the majority of white matter. White matter pathways undergo continuous remodeling during brain maturation (77). Moreover, combined with gene enrichment, previous studies on the development of functional networks, CT, and intracortical myelination have also reported associations with synapse-related terms (17, 23, 78, 79). Our findings may indicate possible synapse-related developmental process mechanisms underlying multiscale structural connectome development from childhood to adolescence.

### Limitations and future directions

There are several limitations to this study. First, our current dataset lacked pubertal hormone measurements, leading us to define ages chronologically instead of by pubertal stage. This limitation may constrain our ability to investigate the effect of pubertal hormone levels on multiscale structural gradients. Incorporating pubertal-related measures into future analyses may yield significant biological insights. Second, the gene expression profiles were exclusively derived from postmortem adult brains, potentially overlooking any developmental impact on gene expression levels. Nevertheless, postnatal spatial gene patterns may exhibit stability (38). To validate our findings, future studies should incorporate pediatric-specific gene expression datasets with spatial resolution comparable to that of the AHBA.

## Materials and Methods

### Participants

We obtained multimodal MR images from the Children School Functions and Brain Development Project in China (Beijing Cohort), which contains a longitudinal dataset of 643 scans from 360 participants (163 females) aged 6-14 years. The final sample included 276 participants (aged 6-14 years, 135 females; 437 scans (159 for 1 timepoint, 83 for 2 timepoints, and 39 for 3 timepoints)) with complete, quality-controlled T1w and T2w images, dMRI scans, and rs-fMRI scans. All participants in this study were cognitively normal, and those with a history of neurological disorders, mental disorders, head injuries, physical illness, or contraindications for MRI were excluded. All study procedures were approved by the Ethics Committee of Beijing Normal University, and written informed consent was obtained from all participants or their parents/guardians.

### Data acquisition

#### MRI acquisition

High-resolution T1w MRI, diffusion MRI, and rs-fMRI data were obtained using 3T Siemens Prisma scanners at Peking University, Beijing, China. T2w scans were acquired using 3T Siemens Prisma scanners at HuiLongGuan Hospital, Beijing, China. The parameters of the T1w scans were as follows: repetition time (TR) = 2530 ms; echo time (TE) = 2.98 ms; inversion time (TI) = 1100 ms; flip angle = 7°; field of view (FOV) = 256 × 224 mm^2^; number of slices = 192; slice thickness = 1 mm; and bandwidth (BW) = 240 Hz/Px. The parameters of the T2w scans were as follows: 3D T2-SPACE sequence, TR = 3200 ms, TE = 564 ms, acquisition matrix = 320 × 320, FOV = 224 × 224 mm^2^, number of slices = 256, slice thickness = 0.7 mm, and BW = 744 Hz/Px. The rs-fMRI scans were acquired using an echo-planar imaging sequence with the following parameters: TR = 2000 ms; TE = 30 ms; flip angle = 90°; FOV =224 × 224 mm^2^; number of slices = 33; number of volumes = 240; and voxel size = 3.5 × 3.5 × 3.5 mm^3^. Diffusion MRI was performed using a high angular resolution diffusion imaging (HARDI) sequence with a 64-channel head coil with the following parameters: TR = 7500 ms, TE = 64 ms, acquisition matrix = 112×112, FOV = 224×224 mm^2^, slices = 70, slice thickness = 2 mm, BW = 2030 Hz/Px, and 64 diffusion weighted directions (b-value = 1000 s/mm^2^) with 10 non-diffusion weighted b0 (0 s/mm^2^).

#### Behavioral data

1. Working memory test. We used a numerical N-back task to estimate WM capacity (48). Twelve blocks of tasks under three workload conditions—0-, 1-, and 2-back—were completed by participants. For the 0-back condition, participants were instructed to judge whether the current digit was 1. For the 1- and 2-back conditions, participants were asked to judge whether the current digit was identical to the previous one or two digits in the sequence. The d-prime index was computed for each condition to assess WM performance. The index was calculated as the inverse of the cumulative Gaussian distribution of the hit ratio subtracted by the inverse of the cumulative Gaussian distribution of the false alarm ratio. The detailed task design can be found in Hao et al.(48). In this study, we included 365 data points.
2. Attentional test. We used a child-friendly version of the Attention Network Test (ANT) (80) to evaluate attention performance, which was measured by the response time for the alerting, orienting and executive control tasks. The detailed task design can be found in Hao et al.(48). We included 372 data points in our study.

### MRI preprocessing

Structural and functional images underwent preprocessing with the modified Human Connectome Project (HCP) pipeline (81).

### Structural MRI

We performed anterior commissure-posterior commissure (AC-PC) alignment and brain extraction. Subsequently, the T1w and T2w images were coregistered using a rigid body transformation with a boundary-based registration cost function (82). Then, the square root of the product of the T1w and T2w images was used to correct for the bias field (83). These images were registered to the Chinese Pediatric Atlas (CHN-PD) (84). Using FreeSurfer 6.0-HCP (85), cortical surfaces were generated in native space, and T2w images were used to refine the pial surfaces. Moreover, cortical ribbon volume myelin maps were generated (83).

### Diffusion MRI

Diffusion images were initially preprocessed using MRtrix3 (86), which included denoising and removing Gibbs ringing artifacts (87). Subsequently, the FSL eddy tool was employed to correct eddy current-induced distortions, head movements, and signal dropout (88–90). Next, the eddy-corrected diffusion images and corresponding field maps were preprocessed using the FSL epi_reg script to effectively mitigate EPI susceptibility artifacts (https://fsl.fmrib.ox.ac.uk/fsl/fslwiki/FLIRT/UserGuide#epi_reg). The diffusion images were finally corrected for B1 field inhomogeneity using the N4 algorithm provided by ANTs (91). Detailed information on the dMRI preprocessing steps can be found in (60).

### Functional MRI

To correct for head motion, each frame of the functional time series was registered to the first frame using rigid body registration. The distortions in the phase encoding direction were corrected using the corresponding field map. The first frame was subsequently registered to the T1w image using rigid body and boundary-based registrations to correct for distortions. The relevant transformations were concatenated to register each frame of functional time series to the first frame, native T1w space, and finally the CHN-PD atlas space. Then, bias field correction, extraction of the brain, and normalization of the whole-brain intensity were performed. Next, followed by a bandpass filter (0.01 Hz < f < 0.08 Hz), we performed ICA-based Automatic Removal Of Motion Artifacts (ICA-AROMA) for denoising (92). We also removed the shared variance between the global signal and time series. Subsequently, the time series in the CHN-PD volume space were projected onto native cortical surfaces using a partial volume weighted ribbon-constrained mapping algorithm. Next, the signals on the cortical surface were resampled and precisely aligned with the Conte69 template through registration, followed by resampling onto the fsaverage5 surface.

### Generation of multiscale structural features

Consistent with the previously reported multiscale model, three complementary structural features were calculated based on T1w, T2w, and diffusion images. The three features were mapped onto Schaefer 1000 parcellations and calculated as described below (the Schaefer 400 atlas was used for validation analysis) (42).

*1) Geodesic distance.* Based on the mid-thickness surface of the individual native surface, the GD was calculated as the shortest distance between two nodes along the surface. In particular, we utilized workbench commands (-surface-geodesic-distance) to compute the distance of each pair of centroid vertices within the given parcel, resulting in a node-by-node GD matrix. Given the limitation of this approach in calculating the GD solely within hemispheres, the interhemispheric GD was calculated by averaging the GD across two hemispheres.
*2) Microstructure profile covariance.* According to a previously reported protocol, we acquired 12 equivolumetric surfaces between the pial and white surfaces and sampled T1w/T2w values along the vertices of these surfaces (6). The intensity profiles of T1w/T2w images were averaged within parcels, excluding any outlier vertices. Then, we calculated pairwise Pearson product-moment correlations between the intensity profiles of each pair of parcels while controlling for the average whole-cortex intensity profile. The matrix was log-transformed after thresholding at zero, resulting in the final MPC matrix.
*3) Tract strength.* We used MRtrix3 to generate a white matter connectivity network. We registered T1w images and their corresponding data to the native diffusion MRI space. An unsupervised algorithm was used to estimate response function (RF) in different brain tissue types (93). Then, we performed single-shell 3-tissue constrained spherical deconvolution (SS3T-CSD) (94) using MRtrix3Tissue (https://3Tissue.github.io), a branch of MRtrix3 (86), to obtain the fiber orientation distribution in all voxels. Following intensity normalization, we chose the gray matter/white matter boundary as the streamline seed mask. Based on anatomically constrained tractography (ACT) (95) with the segmentation results of the structural MR images, second-order integration over fiber orientation distributions was employed to generate streamlines (96). Streamline generation was terminated when 20 million streamlines were counted (maximum tract length = 250 mm; fractional anisotropy cutoff = 0.06; angle threshold = 45°). The spherical deconvolution-informed filtering of tracks (SIFT) approach was used to correct for the bias of streamline density (97). The tract strength (TS) was measured by the number of streamlines. Finally, white matter connectivity was generated by mapping the streamlines onto the Schaefer 1000 atlas and log-transformed.

### Calculation of multiscale structural gradients

We used the BrainSpace Toolbox to compute connectome gradients (https://github.com/MICA-MNI/micaopen/tree/master/structural_manifold) (98). Consistent with a previous study, the nonzero values of the MPC, TS and inverted GD matrices were rank normalized and rescaled to the same numerical range (4). The three matrices were horizontally concatenated and subjected to a diffusion map embedding algorithm with a kernel of normalized angle similarity, which mapped the high-dimensional multiscale structural connectome data into a low-dimensional space (99). The distances in the gradient space reflect dissimilarities in connectivity patterns between regions. In line with previous studies, we set parameter α = 0.5. By dividing the population into 6 groups based on age with 1-year intervals, we generated a group-level multiscale connectome by averaging the individual multiscale matrices. To make the gradients comparable across individuals and eliminate the randomness of the direction of the gradients, we used Procrustes rotations to align the individual gradients to their corresponding age-specific group-level gradients derived from the group-level multiscale connectome (100).

The global gradient measures were computed to summarize the age-related changes in the gradients. These global measures included the following: 1) gradient range, calculated as the difference between the maximum and minimum values; 2) explanation ratio, calculated as the eigenvalue divided by the sum of all eigenvalues; 3) standard deviation, defined as the standard deviation of the given gradient; and 4) gradient dispersion, calculated as the sum of the Euclidean distances of each node to the centroid in the 2D gradient space constructed by the first two gradients. Moreover, we calculated the eccentricity measure as the Euclidean distance between each node and the centroid of the template space obtained from averaging the multiscale matrix across all participants.

### Correlation analysis with cortical morphometric features

To investigate the relationships between multiscale structural gradients and cortical morphometric features, we utilized cortical morphometric features derived from the results of the FreeSurfer preprocessing procedure. Subsequently, 5 cortical morphometric features, CT, GMV, SA, MC, and GC, were extracted and mapped onto the Schaefer 1000 atlas. Given the similarities of cortical patterns across these metrics, we performed PCA to generate a concise representation of the morphometric features. Specifically, for each participant, we conducted PCA on matrix X of node×feature. The first component captured the largest variance, and areas with similar morphological profiles were in close proximity along this principal axis. We conducted a correlation analysis between the first principal component (PC1) and the multiscale structural gradient.

### Calculation of the functional gradient

To assess how structure supported the maturation of functional organization, we related multiscale structural gradients to the FC network. Considering the primary-transmodal functional gradient as a representative of the functional hierarchy and its gradual maturation throughout development, we conducted correlation analysis between structural gradients and functional gradient. We computed pairwise Pearson’s correlation coefficients based on time series with the Schaefer 1000 atlas to obtain individual FC matrices, followed by the generation of a group-averaged FC matrix. We retained the top 10% of edges per row and computed the row-wise normalized angle similarity. This matrix was then input into the diffusion map embedding algorithm, yielding the primary-transmodal functional gradient (99).

### Analysis of multiscale structure–function coupling

We investigated multiscale structure–function coupling during youth, calculated as the Spearman rank correlation between structural connectivity and FC profiles at the nodal level. We computed the average of these individual maps across all participants to generate an averaged coupling map. To quantify the functional specialization of brain networks, we computed the PaC for each scan using the Brain Connectivity Toolbox (https://sites.google.com/site/bctnet/) (101, 102). Based on the Yeo functional networks (45), the PaC measured intermodule connectivity and quantified the extent to which a node participated in other modules.

### Statistical analysis

We employed several LME models to characterize the age effects to adapt for the longitudinal dataset. The candidate models for each measure considered 6 combinations of fixed-effect and random-effect terms, as detailed in Table 1. The mean framewise displacement (mFD) for dMRI was treated as a fixed-effect term and controlled for in this model.

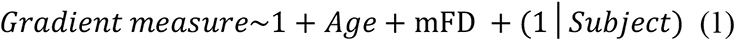

For example, the linear model of the first combination was defined as follows:

We selected the optimal model according to the AIC (46), with a preference for the model exhibiting the lowest AIC value. For regional gradient statistical analysis, we further corrected for multiple comparisons with FDR correction.

For all spatial correlation analyses between different cortical maps, we used the variogram matching approach to estimate the significance (103). By generating 1000 surrogate maps that preserved the spatial autocorrelation of the given brain map, we repeated the correlation analysis utilizing these surrogate maps. The resulting correlation coefficients generated a null distribution comprising 1000 values. The p value was calculated as the proportion of the surrogate coefficients exceeding the actual coefficient.

### Gene enrichment analysis

We collected genome expression data from the AHBA to identify genes associated with age-related multiscale structural gradient changes (https://human.brain-map.org (50)). The AHBA is a regional microarray transcriptomic dataset of 3702 tissue samples from 6 healthy adult donors. We used the abagen toolbox (version 0.1.3; https://github.com/rmarkello/abagen) to preprocess the microarray data using the Schaefer 1000 atlas. Given that right hemisphere data were only available from 2 donors, we opted to utilize the data from the left hemisphere for our analysis. Using the default parameters, we finally obtained a 416 ×15631 (region × gene) matrix.

To determine the relationships between age-related changes in the multiscale structural gradient and genes, we used the previously obtained age effect t statistics of the principal gradient (t-map) and gene expression matrix in partial least squares (PLS) regression. Our goal was to identify the components associated with the gradient t-map, which represented optimally weighted linear combinations of expression patterns. The first component (PLS1) was the most strongly correlated with the t-map. By using a previously described spatial autocorrelation correction approach, we examined the statistical significance of the variance explained by the PLS components and the correlation coefficient between PLS1 and the t-map (103). Subsequently, bootstrapping was performed to assess the error of each gene weight from PLS1, and we transformed the weights into Z scores by dividing the weight by the standard deviation of the given weight derived from 1000 bootstrapping results. We selected the top 10% of genes from both the positive and negative weights, which made the largest contribution to PLS1, for the subsequent gene enrichment analysis.

The positive and negative genes were then separately entered into the Metascape webtool for gene enrichment analysis (51). According to GO analysis, Metascape was used to search for specific molecular function, biological process, and cellular component terms. The resulting enriched pathways were thresholded for significance at an FDR < 5%.

### Analysis of the relationship between cognition and the principal multiscale structural gradient

We performed PLSC analysis (104) with the myPLS toolbox (https://github.com/danizoeller/myPLS) to extract the relationships between the multiscale structural gradient and cognitive scores. PLSC analysis was performed separately for WM and attention performance. We first computed a covariance matrix R between brain variables X and cognition variables Y:

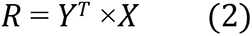

followed by singular value decomposition on R:

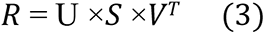

where U and V reflect the contributions of the cognition and brain variables, respectively, to the LCs, while S represents the singular values. Then, brain scores (Lx = X×V) and cognition scores (Ly=Y×U) were computed for each LC by projecting brain and cognition variables onto their corresponding weights. Brain loadings and cognition loadings were computed as Pearson correlations between the original data and previously obtained scores. Overall, the PLSC analysis generated LCs that represented the optimal weighted linear combinations of the original variables, thereby establishing the strongest relationships between the brain and cognition data. Subsequently, we assessed the statistical significance of each LC using a permutation test (n=1000). Specifically, we randomly shuffled the cognitive data across all subjects, resulting in a null distribution of singular values. By comparing the actual value with the null distribution, we ascertained the statistical significance. The statistical significance of brain and cognition loadings was estimated by bootstrap resampling (n=1000), with replacement across all subjects on X and Y.

## Declaration of Competing Interests

The authors declare no competing interests.

## Acknowledgments

The authors would like to thank all the families and children for their support and participation.

## Abbreviations

AIC: Akaike information criterion
AHBA: Allen Human Brain Atlas
ANT: Attention Network Test
CBD: Children School Functions and Brain Development Project in China (Beijing Cohort)
DMN: default mode network
DAN: dorsal attention network
EC: executive control
FPN: frontoparietal network
FC: functional connectivity
FDR: false discovery rate
GD: geodesic distance
GC: Gaussian curvature
GO: Gene Ontology
HARDI: high angular resolution diffusion imaging
HCP: Human Connectome Project
LN: limbic network
LME: linear mixed-effect
LC: latent component
MPC: microstructural profile covariance
MC: mean curvature
PCA: principal component analysis
PLSC: partial least square correlation
PLSR: partial least squares regression
PaC: participation coefficient
S-A: sensorimotor-association
SA: surface area
SN: somatomotor network
TS: tract strength
VAN: ventral attention network
WM: working memory.

## Supporting information

**S1 Fig. The first two multiscale structural gradients projected onto the cortical surface for each group.**

**S2 Fig. Developmental pattern of the second multiscale structural gradient. (A)** Radar plot of the second gradient for comparison between 6-7 years group and other groups based on Yeo functional networks (left) (45) and laminar differentiation parcellation (right) (44). (**B)** Global density map of the second gradient for each group. **(C)** Correlation coefficient between the second structural gradient and primary-to-transmodal functional gradient changed across age (not significant).

**S3 Fig. Multiscale structural gradients during childhood and adolescence based on Schaefer 400 atlas. (A)** The group-level gradients based on the Schaefer 400 atlas exhibited a spatial pattern that was highly consistent with those derived from the Schaefer 1000 atlas. (**B)** Global density map of the first two gradients for each group showed a similar pattern with those derived from the Schaefer 1000 atlas. (**C)** Radar plot of the first two gradients for comparison between 6-7 years group and other groups based on Yeo functional networks (45). **(D)** The first two structural gradients mapped into a 2D gradient space for 6-7, 9, and 12-13 years old group demonstrated an expansion pattern during development.

**S4 Fig. Spatial correlation between multiscale structure-function coupling map and primary-to-transmodal functional gradient (A) as well as multiscale structural gradient (B) (p _surrogate_<0.01).**

**S5 Fig. Age-related changes in multiscale structure-function coupling.** Age-related increases/decreases were shown in red/blue. The right panel showed t-values distribution based on Yeo functional networks.

**S6 Fig. Association between age-related changes in principal gradient and gene expression profiles. (A)** Gene Ontology (GO) enrichment pathways of top 10% genes with positive PLS 1 weights. The most significant 20 GO terms were displayed (left panel). Metascape enrichment network visualization showed the intra-cluster and inter-cluster similarities of enriched terms. **(B)** Gene Ontology (GO) enrichment pathways of top 10% genes with negative PLS 1 weights.

**S1 Movie. The developmental process of the gradient space across different ages in Schaefer 1000 atlas.**

**S2 Movie. The developmental process of the gradient space across different ages in Schaefer 400 atlas.**

## Notes

### Competing Interest Statement

The authors have declared no competing interest.

